# A Synthetic Microbial Operational Amplifier

**DOI:** 10.1101/161828

**Authors:** Ji Zeng, Jaewook Kim, Areen Banerjee, Rahul Sarpeshkar

## Abstract

Synthetic biology has created oscillators, latches, logic gates, logarithmically linear circuits, and load drivers that have electronic analogs in living cells. The ubiquitous operational amplifier, which allows circuits to operate robustly and precisely has not been built with bio-molecular parts. As in electronics, a biological operational-amplifier could greatly improve the predictability of circuits despite noise and variability, a problem that all cellular circuits face. Here, we show how to create a synthetic 3-stage inducer-input operational amplifier with a differential transcription-factor stage, a CRISPR-based push-pull stage, and an enzymatic output stage with just 5 proteins including dCas9. Our ‘Bio-OpAmp’ expands the toolkit of fundamental circuits available to bioengineers or biologists, and may shed insight into biological systems that require robust and precise molecular homeostasis and regulation.

**One Sentence Summary:** A synthetic bio-molecular operational amplifier that can enable robust, precise, and programmable homeostasis and regulation in living cells with just 5 protein parts is described.

---

Several electronic circuit motifs and devices including oscillators(*1–3*), latches (*4, 5*), logic gates(*6–8*), logarithmically linear circuits (*9*), and load drivers (*10*) have been designed and ported to biological systems and applications. However, the operational amplifier, which is a fundamental and highly useful electronic device in negative-feedback and regulatory loops, has never been ported to biology. Negative-feedback and regulatory loops are common in biology (*11*) and in electronics. In both cases, the use of negative feedback can provide seven benefits including insensitivity to parameters, rejection of disturbing inputs, linearization of nonlinear components, impedance or buffering transformations, speedup, implementation of inverse functions, and stabilization of unstable systems (*12*).

Operational amplifiers implement a simple circuit building block wherein a strong driving output is a highly amplified version of the difference of two of its control inputs. They enable straightforward implementations of negative-feedback analog circuit motifs (*12–14*) with high precision and large robustness. The operational amplifier has been incredibly important in making analog electronics and analog computation scalable, technology, and context independent from an input-output point of view. Hence, operational amplifiers are as ubiquitous in analog circuits and analog computation as logic gates are in digital circuits. Not surprisingly, currently nearly 10M operational amplifiers are sold every day.

Ideas from analog circuits and feedback theory are increasingly being appreciated as very important in synthetic and systems biology (*11, 13–24*). We discuss two examples that are useful for providing some motivation for our work.

A quorum-sensing molecule has been used to increase the production of a cell killing molecule such that attenuations in the fluctuations of a microbial population result (*20*). Such a circuit is an example of a clever negative-feedback loop that achieves some population control in living cells. However, it does not use an operational amplifier with a user-programmable population set-point to achieve precise population control. Thus, the circuit would not be robust to disturbances in the quorum sensing molecule concentration. As another example, prior ‘load driver’ circuits in cells have used an elegant phosphor-relay system to create large production and degradation fluxes at their outputs, which are inherently fast, such that relatively small loading fluxes, which are inherently slow, do not significantly impact their operation (*10*). Thus, a ‘molecular buffer circuit’ can be created. However, it does not use an operational amplifier with global negative feedback around all its gain stages to achieve high feedback loop strength. Thus, unlike true operational-amplifier topologies, it does not allow for robust closed-loop programmable gain, is not insensitive to nonlinearities from its input to output, and cannot provide good buffering capability at low output flux.

Figures 1a, 1b, and 1c illustrate our three-gain-stage biological operational amplifier (Bio-OpAmp) as a conceptual block diagram, as it is implemented in cells with molecular parts, and as a circuit schematic respectively. The first gain stage is a fast-differential stage with arabinose and AHL inducer inputs that generate two corresponding *sgRNA* outputs. The second gain stage is a slow differential stage that uses CRISPR-*sgRNA* complexes to produce AiiA and LuxI enzyme outputs. The third gain stage is a fast stage that uses AiiA and LuxI enzymes to help degrade or increase AHL concentration respectively (*2*). The AiiA and LuxI enzymes also perform differential-to-single-input conversion to regenerate the AHL input as an output such that a negative feedback loop is created. In this negative-feedback loop, arabinose then effectively serves as a control or set-point inducer input that determines the intended equilibrium and homeostatically-controlled value of the output inducer AHL. Since protein (green) production is relatively slow, but RNA (red) and inducer (purple) production are fast, the negative-feedback loop inherently has a dominantly slow protein time constant corresponding to LuxI/AiiA degradation. This dominant time constant naturally promotes stability even under strong negative feedback (*12*), much like that of a dominant integrator time constant in well-known operational-amplifier or biological feedback configurations in chemotaxis (*17*).

**Fig. 1.**
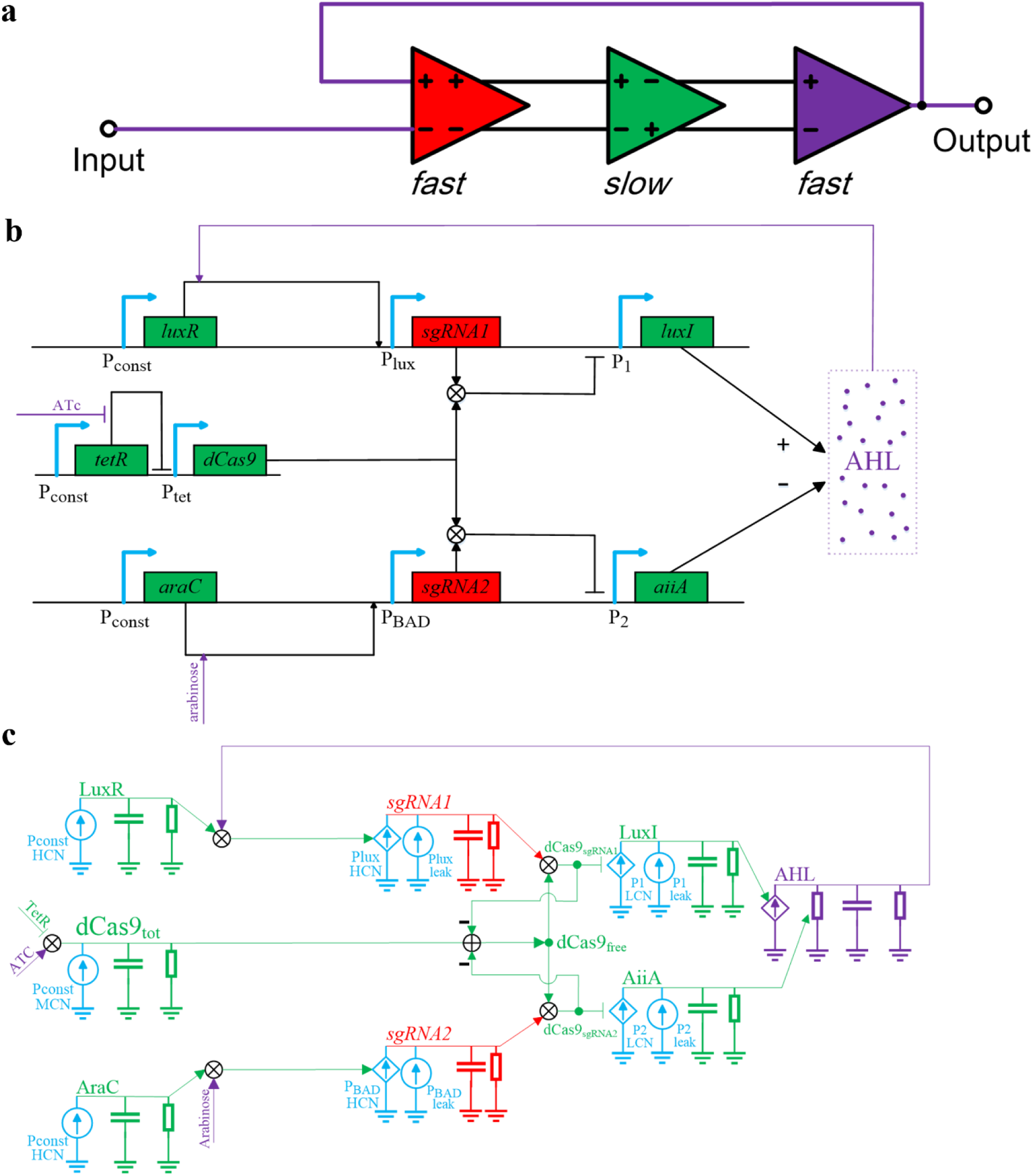
Circuits of the biological operational amplifier. **a**. A block diagram of a three-gain-stage amplifier (OpAmp) with negative feedback for robust closed-loop operation. **b.** The circuit for a biological operational amplifier (Bio-OpAmp). LuxR and AraC are produced from constitutive promoters. In addition, dCas9 concentration is controlled by ATc. The red-colored *sgRNA* and purple-colored inducer AHL have fast dynamics while green-colored proteins have faster dynamics. **c.** An analog circuit schematic for the Bio-OpAmp. Such a schematic can quantitatively represent the differential equations of the Bio-OpAmp dynamical system (*14*) and is useful for porting existing electronic design, modeling, and simulation tools to biological design, modeling, and simulation (see Supplementary and **Figure 3**). Four basic kinds of molecular circuit elements are represented with different colors: DNA promoters (blue), RNA (red), protein (green), and small–molecule inducer (purple). Production of species are represented by dependent (arrow-controlled diamond generators) or independent (circular) current generators, degradation of species by resistors (rectangular elements), and preservation of molecular concentration state by capacitors (horizontal parallel-line elements).

The CRISPR-based parts in Figure 1b were constructed by modifying a plasmid used in other circuits (*25*). The CRISPR system was selected mainly because dCas9 can be programmed to bind almost any chromosome locus with a PAM recognition site with high specificity and because RNA dynamics are fast. Even though the feedback loop strength is determined by three gain stages and is therefore relatively large, the short life time of *sgRNA (26)* and AHL in our system compared with the long-life time of LuxI and AiiA naturally alleviate oscillation and instability. Such multi-gain-stage amplifiers, are otherwise, prime candidates for oscillation (*12*) as for example in repressor cascades with negative feedback (*1*). Figure 1c is useful in mapping our biological circuit schematic to electronic circuit schematics that quantitatively and pictorially model and represent the production (diamond current-producing generators), degradation (rectangular current-consuming resistors) or state preservation (horizontal-parallel-line capacitors) of molecular species in coupled differential equations with DNA (blue), RNA (red), protein (green), or small-molecule inducers (purple) (*14*). Current generation at DNA promoters is either constitutive (circular current generator) or dependent on another control variable (diamond current generator with an arrow input). Figure 1c reveals that because, dCas9 is a fixed resource, whose molecule number is regulated by ATc/TetR, its binding or consumption by one half of the circuit, i.e., by *sgRNA1*, leads to less of it being available to bind to the other half of the circuit, i.e., by *sgRNA2*. Thus, there is fast coupling between the two halves of the operational amplifier that effectively serves to increase its gain. Others have also exploited the sharing of a common resource to benefit their synthetic circuits via fast coupling as well (*2*). In our case, the sharing of dCas9 in the Bio-OpAmp effectively serves to increase gain via ‘push-pull’ amplification. A push-pull effect is exploited in the output stages of many electronic operational amplifiers to improve their performance as well (*12*).

Figures 2a and 2c show each differential half of the first two gain stages of the Bio-OpAmp of Figure 1 with the operational amplifier operated in open-loop fashion (without negative feedback). These halves were built by simply not implementing the other half in plasmids that we tested separately. Figures 2b and 2d show corresponding measured data in *E. coli*. Both of our circuits, P_1_-*rfp* and P_2_-*rfp*, were implemented as three-plasmid systems. In the high copy number plasmid (HCN), *sgRNA1* is controlled by a constitutively expressed LuxR (*27*) while *sgRNA2* is controlled by AraC (*28*) respectively. The dCas9 shared-resource pool is controlled by TetR (*29*) and ATc (anhydrotetracycline).

**Fig. 2.**
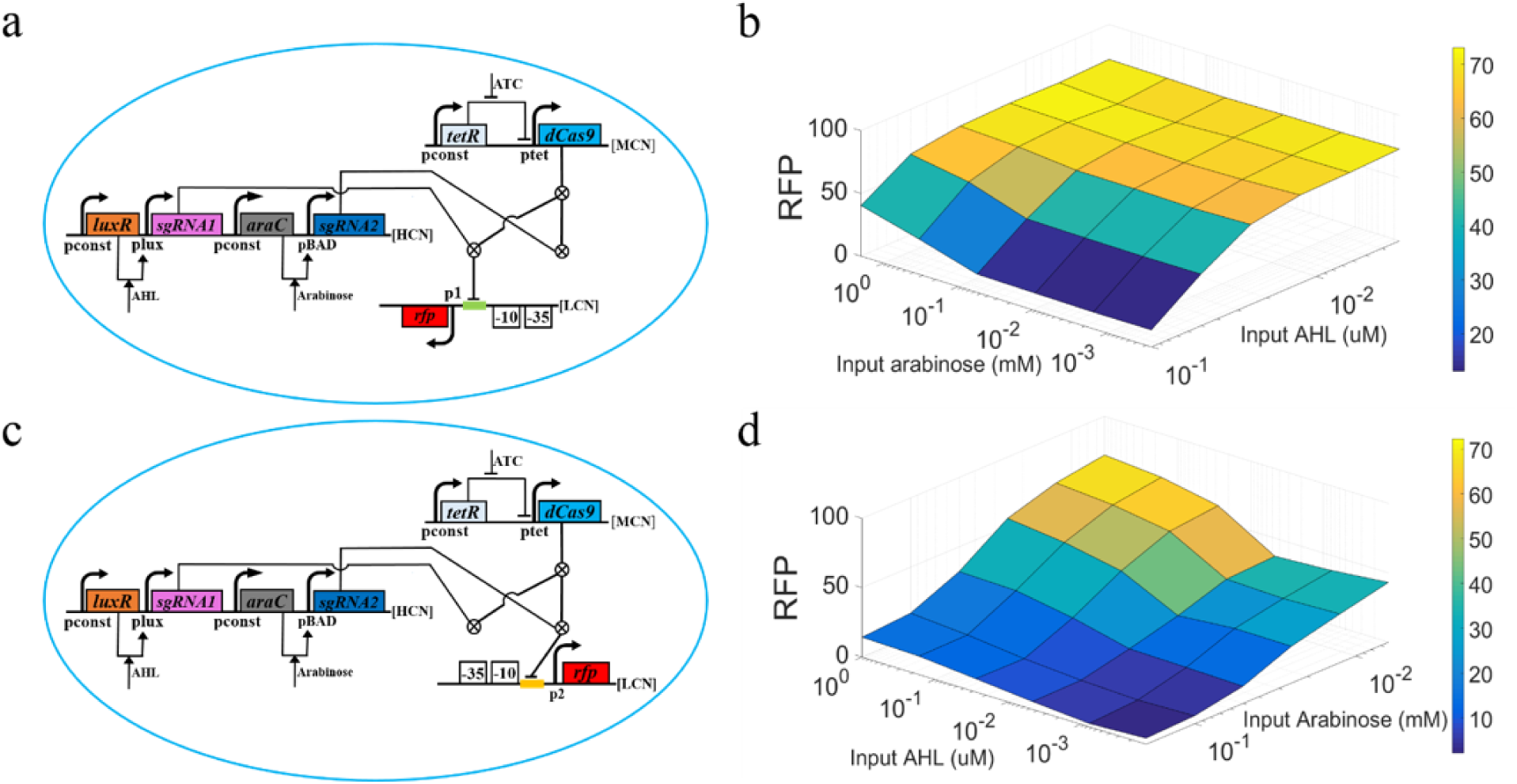
Experimental characterization of the two half circuits of the biological operational amplifier under open-loop operation. P_1_*-rfp* half circuit of the operational amplifier. *sgRNA1* and *sgRNA2* compete for dCas9 binding. The *sgRNA1* binding site is placed between a -10 site and a transcriptional start site to create synthetic repression at the *p*_*1*_ promoter. An *rfp* gene is ligated to P1 in the low copy number plasmid (LCN). **b.** The push and pull effects in P_1_*-rfp* caused by the shared dCas9 resource between *sgRNA1* and *sgRNA2*. X and Y axes indicate the input concentration of AHL and arabinose and the Z axis represents the output RFP level. AHL represses P_1_*-rfp*, while arabinose promotes de-repression when its concentration is high enough to significantly ‘decoy’ dCas9. **c.** The P_2_-*rfp* half circuit of the operational amplifier. An *sgRNA2* binding site architects’ repression of the synthetic P_2_ promoter. **d.** The push and pull effects in P_2_*-rfp* caused by the shared dCas9 resource. Here, arabinose represses P_2_*-rfp*, while AHL de-represses it analogous to the effects described in **a** and **b**.

Even though arabinose does not effectively connect to the repressed output in Figure 2a as AHL does, at relatively high concentrations, it competes for dCas9 as a ‘decoy’. Thus, it leads to de-repression and increase of the AHL-controlled RFP output in Figure 2b, a manifestation of the push-pull shared-resource effect of Figure 1c. Similarly, even though AHL does not effectively connect to the repressed output in Figure 2b as arabinose does, at relatively high concentrations, it competes for dCas9 as a ‘decoy’. Thus, it leads to de-repression and increase of the arabinose-controlled RFP output in Figure 2d. Due to the asymmetric leaky expression of Plux and PBAD, or the asymmetric binding Kd’s of *sgRNA*s to dCas9, or the resulting complexes to the DNA promoters, the push-pull effects of P_1_-*rfp* and P_2_-*rfp* in Figures 2a and 2c respectively are not symmetric (Figures 2b and 2d). Nevertheless, as the biological data of Figure 3 show, just as in electronics, if there is strong negative feedback in a closed-loop operational amplifier, such effects only serve to vary the net open-loop gain of the operational amplifier at different operating points and have relatively little impact on its robust closed-loop performance.

**Fig. 3.**
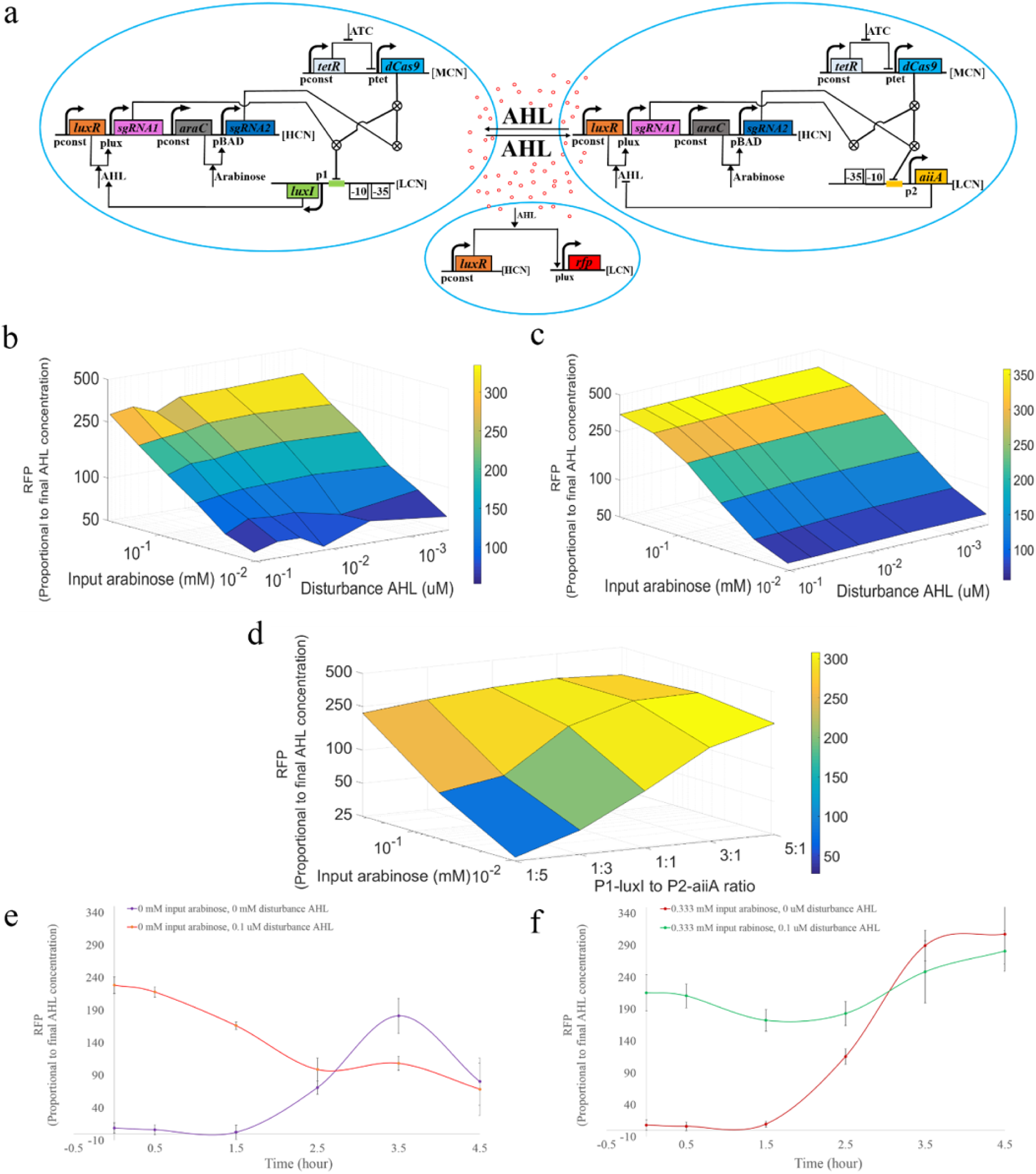
Experimental characterization of the full biological operational amplifier under closed-loop operation. **a**, The circuit schematic of the complete biological operational amplifier. AHL is produced by P_1_*-luxI* and degraded by P_2_*-aiiA*. The microbial consortia communicate with each other via AHL. The AHL concentration is measured by Plux*-rfp* circuit (see supplementary Figure 1) such that RFP acts as a proxy for the final output AHL. **b**. Input-output characteristics of the biological operational amplifier. The output RFP only tracks controlling input arabinose while rejecting an externally added initial AHL input disturbance. **c.** Circuit simulation data of the biological operational amplifier represented and modeled by the schematic of Figure 1c. The data are in good agreement with the experimentally measured data shown in **b**. Electronic circuit design tools (CADENCE) were used to create and simulate the biological operational amplifier. **d**. Varying closed-loop gain in the biological operational amplifier. Different ratios of P_1_*-luxI* and P_2_*-aiiA* circuits lead to different closed-loop gains (variation in RFP with varying arabinose input) in the biological operational amplifier. **e** and **f**. Stable dynamics towards the arabinose-determined set-point in the closed-loop operational amplifier. The experimentally measured data reveal that, in all cases, the final outputs of the closed-loop biological operational amplifier equilibrate very near a value determined by the arabinose setpoint regardless of the value of the disturbance AHL input. Since the arabinose controlling input is higher in **f** than in **e,** the final equilibrium output is higher in **f** than in **e.**

Figure 3a illustrates circuits and measurements from the full three-gain-stage closed-loop Bio-OpAmp with input arabinose and output AHL. We replaced *rfp* with *luxI* in the P_1_-*luxI* circuit of Figure 2a and *rfp* with *aiiA* in the P_2_-*aiiA* circuit of Figure 2c to create the circuit of Figure 3. The output AHL was measured via the proxy *rfp* molecule as shown (Supplementary Figure S1). We tested the full Bio-OpAmp by combining strains with P_1_*-luxI* circuits and strains with P_2_*-aiiA* circuits in a common co culture (details are described in materials and methods). The closed-loop amplification can then be programmably and conveniently varied by varying the cell numbers in these two populations. It also alleviates synthetic-circuit metabolic burden in each cell as in several other practicable synthetic-circuit schemes (*2, 3, 8, 30*). Finally, it could enable a modular design for our Bio-OpAmp that can be ported by others to cellular circuits with different consortia and parts in a molecular-homeostasis application of their choosing. Supplementary Table S2 suggests a few examples.

Measured experimental data in Figure 3b reveal that the output AHL faithfully tracks the input arabinose over more than an order of magnitude of input dynamic range as would be expected in a high-performance control system with negative feedback. Furthermore, Figure 3b also shows that the operational amplifier is robust to intentionally induced initial disturbance inputs of AHL over more than two orders of magnitude, always rejecting these disturbances such that the final measured AHL output (or its proxy *rfp*) only tracks the arabinose input. Figure 3c shows that a model of the Bio-OpAmp based on Figure 1c, which is described in more detail in the Supplementary section, yields results very near that measured experimentally in Figure 3b. Supplementary Figure 2 shows that, in a control experiment where there is no synthetic negative-feedback operational amplifier present, the disturbance AHL input is not rejected such that the final AHL output concentration is simply proportional to the disturbance AHL concentration, and does not track the arabinose input.

Figure 3d shows that the gain (output *rfp* variation for a given variation in arabinose) varies when the ratio of P_1_*-luxI* to P_2_*-aiiA* varies from 1:5 to 5:1 in the Bio-OpAmp such that user-programmable closed-loop gain can be implemented. In our case, a nearly 10-fold change in output results for low arabinose concentrations and different plasmid (or cell-population) ratios. Logarithmically linear operation in Figure 3 arises because of the logarithmically linear biochemical potential variation in analog circuits and analog computation in cells (*9, 12–14*).

Figures 3e and 3f show that, for two different arabinose input conditions and different disturbance AHL inputs, the final equilibrium of the closed-loop amplifier dynamically approaches that given by the arabinose input set-point and completely rejects the disturbance AHL input. Also, as expected from the circuit of Figures 1 or 3, higher arabinose input leads to higher cellular output. The dynamics of the closed-loop operational amplifier does not show strong overshoots, undershoots, or oscillation indicating that the negative-feedback loop is stable. It is worth noting that the data of Figures 3e and 3f would not show good input-output behavior if the closed-loop operational amplifier were unstable.

Homeostasis is a universal phenomenon wherein biological systems maintain steady states of output variables rejecting disturbances from external conditions (*11, 17, 19*). Bacterial iron homeostasis, plasma-ionized calcium homeostasis, and human blood glucose homeostasis provide but a few examples (*21–23*). In biology, just as in our synthetic circuit, two different molecules compete via two different halves of a negative-feedback loop (*21–23*) to arrange molecular homeostasis. For example, blood glucose homeostasis in the body, which malfunctions in diabetes, is implemented through two half circuits related to insulin-releasing β cells in the pancreas and glucagon-releasing α cells in the pancreas (*21, 22*). Hence, our synthetic circuit with the AHL and arabinose halves has similar analogs in nature.

Our Bio-OpAmp circuit motif may be useful in molecular homeostasis in biotechnology and medicine. In Table S2 in the supplementary, we suggest how Bio-OpAmps in different cell types can be designed with different molecular parts pertinent to the cell or application but with the same circuit architecture. For example, in technologies that utilize microbes to produce industrially important molecules like succinic-acid, toxicity or substrate inhibition from intermediate metabolites or final products is one of the reasons that bacterial growth is inhibited (*24, 31*). A Bio-OpAmp can enable precise control of such metabolites.

Over nearly two decades, others have shown that logic-gate circuit motifs can be ported across various cell types using different parts but with the same logic circuit architecture. Similarly, in a future medical application, a change of the 4 proteins involved in stages 1 and 3 of our circuit motif, i.e., new small-molecule inputs and new push-pull enzymes in stages 1 and 3 with dCas9-RNA complexes altered for therapeutic promoters of interest, may enable repair of natural feedback loops involved in homeostatic dysfunction. Finally, schematic circuit motifs like those in Figure 1c that rigorously represent and model biological circuits like those in Figure 1b and general methodologies for generating them (*14*) can help port a large class of analog electronic circuit topologies to biological equivalents. (*32–40*)

## Acknowledgments

We thank Alec A. K. Nielsen, a member of Christopher A. Voigt lab, for providing plasmids pAN-L3S3P21-PTet-dCas9 and pAN-NOR, which formed a good initial starting point for our dCas9 CRISPR constructs. These plasmids were substantially modified by us for this work. This work was supported by funds to R. Sarpeshkar from Dartmouth College and M.I.T.

## Supplementary Materials

Materials and Methods

Supplementary Text

Tables S1 & S2

Figures S1 & S2

## References and Notes

1. M. B. Elowitz, S. Leibler, A synthetic oscillatory network of transcriptional regulators. Nature 403, 335–338 (2000).

2. A. Prindle et al., Rapid and tunable post-translational coupling of genetic circuits. Nature 508, 387–391 (2014).

3. Y. Chen, J. K. Kim, A. J. Hirning, K. Josic, M. R. Bennett, SYNTHETIC BIOLOGY. Emergent genetic oscillations in a synthetic microbial consortium. Science 349, 986–989 (2015).

4. T. S. Gardner, C. R. Cantor, J. J. Collins, Construction of a genetic toggle switch in Escherichia coli. Nature 403, 339–342 (2000).

5. J. W. Lee et al., Creating Single-Copy Genetic Circuits. Mol Cell 63, 329–336 (2016).

6. M. N. Win, C. D. Smolke, Higher-order cellular information processing with synthetic RNA devices. Science 322, 456–460 (2008).

7. J. Bonnet, P. Yin, M. E. Ortiz, P. Subsoontorn, D. Endy, Amplifying genetic logic gates. Science 340, 599–603 (2013).

8. A. A. Nielsen et al., Genetic circuit design automation. Science 352, aac7341 (2016).

9. R. Daniel, J. R. Rubens, R. Sarpeshkar, T. K. Lu, Synthetic analog computation in living cells. Nature 497, 619–623 (2013).

10. D. Mishra, P. M. Rivera, A. Lin, D. Del Vecchio, R. Weiss, A load driver device for engineering modularity in biological networks. Nat Biotechnol 32, 1268–1275 (2014).

11. U. Alon, An Introduction to Systems Biology. (Chapman & Hall/CRC, 2016).

12. R. Sarpeshkar, Ultra Low Bioelectronics: Fundamentals, Biomedical Applications, and Bio-inspired systems. (Cambridge University Press, 2010).

13. R. Sarpeshkar, Analog synthetic biology. Philosophical Transactions of the Royal Society a-Mathematical Physical and Engineering Sciences 372, (2014).

14. J. J. Y. Teo, S. S. Woo, R. Sarpeshkar, Synthetic Biology: A Unifying View and Review Using Analog Circuits. Ieee Transactions on Biomedical Circuits and Systems 9, 453–474 (2015).

15. H. M. Sauro, K. H. Kim, Synthetic biology: It′s an analog world. Nature 497, 572–573 (2013).

16. T. Song, S. Garg, R. Mokhtar, H. Bui, J. Reif, Analog Computation by DNA Strand Displacement Circuits. ACS Synth Biol 5, 898–912 (2016).

17. T. M. Yi, Y. Huang, M. I. Simon, J. Doyle, Robust perfect adaptation in bacterial chemotaxis through integral feedback control. Proc Natl Acad Sci U S A 97, 4649–4653 (2000).

18. P. E. M. Purnick, R. Weiss, The second wave of synthetic biology: from modules to systems. Nature Reviews Molecular Cell Biology 10, 410–422 (2009).

19. D. D. Vecchio, R. M. Murray, Biomolecular Feedback Systems. (Princeton University Press, 2014).

20. L. You, R. S. Cox, 3rd, R. Weiss, F. H. Arnold, Programmed population control by cell-cell communication and regulated killing. Nature 428, 868–871 (2004).

21. G. Jiang, B. B. Zhang, Glucagon and regulation of glucose metabolism. Am J Physiol Endocrinol Metab 284, E671–678 (2003).

22. I. Quesada, E. Tuduri, C. Ripoll, A. Nadal, Physiology of the pancreatic alpha-cell and glucagon secretion: role in glucose homeostasis and diabetes. J Endocrinol 199, 5–19 (2008).

23. G. R. Mundy, T. A. Guise, Hormonal control of calcium homeostasis. Clin Chem 45, 1347–1352 (1999).

24. V. Chubukov, A. Mukhopadhyay, C. J. Petzold, J. D. Keasling, H. G. Martín, Synthetic and systems biology for microbial production of commodity chemicals. Npj Systems Biology And Applications 2, 16009 (2016).

25. A. A. Nielsen, C. A. Voigt, Multi-input CRISPR/Cas genetic circuits that interface host regulatory networks. Mol Syst Biol 10, 763 (2014).

26. H. Ma et al., CRISPR-Cas9 nuclear dynamics and target recognition in living cells. J Cell Biol 214, 529–537 (2016).

27. D. M. Sitnikov, J. B. Schineller, T. O. Baldwin, Transcriptional regulation of bioluminesence genes from Vibrio fischeri. Mol Microbiol 17, 801–812 (1995).

28. R. Schleif, AraC protein, regulation of the l-arabinose operon in Escherichia coli, and the light switch mechanism of AraC action. FEMS Microbiol Rev 34, 779–796 (2010).

29. J. L. Ramos et al., The TetR family of transcriptional repressors. Microbiol Mol Biol Rev 69, 326–356 (2005).

30. S. G. Hays, W. G. Patrick, M. Ziesack, N. Oxman, P. A. Silver, Better together: engineering and application of microbial symbioses. Curr Opin Biotechnol 36, 40–49 (2015).

31. T. Warnecke, R. T. Gill, Organic acid toxicity, tolerance, and production in Escherichia coli biorefining applications. Microb Cell Fact 4, 25 (2005).

32. T. Danino, O. Mondragon-Palomino, L. Tsimring, J. Hasty, A synchronized quorum of genetic clocks. Nature 463, 326–330 (2010).

33. H. M. Sauro, Enzyme Kinetics for Systems Biology. (Ambrosius Publishing, 2011).

34. P. Wang, G. Bashiri, X. Gao, M. R. Sawaya, Y. Tang, Uncovering the enzymes that catalyze the final steps in oxytetracycline biosynthesis. J Am Chem Soc 135, 7138–7141 (2013).

35. X. Wen, Y. Jia, J. Li, Enzymatic degradation of tetracycline and oxytetracycline by crude manganese peroxidase prepared from Phanerochaete chrysosporium. J Hazard Mater 177, 924–928 (2010).

36. S. Patel, Disruption of aromatase homeostasis as the cause of a multiplicity of ailments: A comprehensive review. J Steroid Biochem Mol Biol, (2017).

37. M. Schicht et al., Articular cartilage chondrocytes express aromatase and use enzymes involved in estrogen metabolism. Arthritis Res Ther 16, R93 (2014).

38. K. K. Cheng, G. Y. Wang, J. Zeng, J. A. Zhang, Improved succinate production by metabolic engineering. Biomed Res Int 2013, 538790 (2013).

39. M. H. Kim et al., The molecular structure and catalytic mechanism of a quorum-quenching N-acyl-L-homoserine lactone hydrolase. Proc Natl Acad Sci U S A 102, 17606–17611 (2005).

40. A. L. Schaefer, D. L. Val, B. L. Hanzelka, J. E. Cronan, Jr., E. P. Greenberg, Generation of cell-to-cell signals in quorum sensing: acyl homoserine lactone synthase activity of a purified Vibrio fischeri LuxI protein. Proc Natl Acad Sci U S A 93, 9505–9509 (1996).

